# Human Haematopoietic Stem Cells released into circulation following mobilization express multiple G0-associated quiescence markers

**DOI:** 10.64898/2026.06.15.732175

**Authors:** Jos Domen, Rahul Sinha, Daniel Dan Liu, Benjamin Ohene-Gambill, Jason B. Ross, Norma F. Neff, Irving L Weissman

**Affiliations:** Institute for Stem Cell Biology and Regenerative Medicine, School of Medicine, Stanford University, Stanford, CA 94305; Ludwig Cancer Center at Stanford, Stanford University, Stanford, CA 94305; Department of Pathology, Stanford University, Stanford, CA 94305; Chan Zuckerberg Biohub SF, San Francisco CA 94158, USA; 930 Lundy Lane, Los Altos, CA94024; Dept of Radiation Oncology, Stanford University, Stanford, CA94305; Stanford Cancer Institute, Stanford University School of Medicine, Stanford CA94305

## Abstract

Haematopoietic stem cells (HSC), while usually quiescent, can rapidly divide following specific stimuli (mobilization). These HSC can seed additional niches, allowing for the swift generation of essential blood cells. However, studies in mice and humans have clearly demonstrated that cycling bone marrow (BM) HSC (cells in the G1/S/G2/M phases) engraft and reconstitute the haematopoietic system poorly compared with HSC in the G0 phase^1^. This raises the question why mobilized HSC, immediately following 3 or more cell divisions^2^, efficiently reconstitute the haematopoietic system. We studied this phenomenon in human HSC using scRNAseq analysis. We found that mobilized HSC rapidly start transcribing genes associated with quiescence, specific for the G0 phase of the cell cycle. We hypothesize that this rapid switch from actively dividing to quiescent cells combined with our extensive RNA expression data will allow us to better define pathways involved in this process.

## INTRODUCTION

HSC travel to different organs during development, from yolk sac to fetal liver to spleen. Most settle in the BM, some can be found in other organs^3^. Adult HSC can exit the BM and travel to spleen, liver and other organs^4^. This may be a defensive mechanism to ensure maintenance of HSC, e.g. following exposure to toxins or severe blood loss.

Mobilization typically involves two phases, (*i*) proliferation, (*ii*) release into circulation. Growth factors and chemotherapeutics induce cell divisions and rapid expansion in the BM. Disruption of retention signals (CXCR4/CXCL12, VLA-4/VCAM-1) allows egress of the expanded HSC population, as mostly non-cycling cells^5^. Newer mobilizers, including AMD3100/Plerixafor, a CXCR4 antagonist, can induce rapid mobilization in a proliferation independent fashion^6,7^.

Preclinically^8^, and clinically^9^, this mechanism is used to harvest HSC-containing cell preparations through apheresis, a process that has, for the most part, replaced BM harvest. Several methods have been developed, using growth factors (e.g. G-CSF), often combined with chemotherapeutics such as cyclophosphamide^10^,^11^. Mouse estimates show that the HSC need to undergo 3-4 divisions in 3 days to produce sufficient HSC^8^. Optimization of human mobilization protocols remains under active investigation^9,12^.

HSC mobilization has revealed an unexpected and counterintuitive cell cycle aspect. It has been well established that the relatively rare, BM-derived, actively cycling HSC (G1/S/G2M) are engraft poorly^13^. HSC that are not actively cycling (G0) engraft efficiently. Mobilized Peripheral Blood (MPB) HSC, have spent 3-4 days actively and rapidly cycling, before egressing and rapidly (<30 seconds) transiting through the blood, mostly to spleen and liver and a lesser extent, BM^4,11^. HSC that cycle immediately prior to release into the bloodstream, engraft efficiently. HSC released into the blood during a mobilization protocol have few cells in S/G2/M, most are diploid^11^. Engraftment potential of G0 HSC is robust and sustained, while G1 HSC engraftment is much less so^1^. Yet MPB stem cells, directly after several rapid cell divisions that must have come through G1 to enter the next S phase, engraft efficiently. This raises a question: Are these cells in G1 or G0?

To address this we studied human HSC, mobilized using either G-CSF or G-CSF plus plerixafor. HSC are defined as singlets that are PI-Lin-CD34+ CD38-CD90+ CD45RA-. Haematopoietic multi-potent Progenitor Cells (both MPP and LMPP), differ from HSC in that they are CD90-. Single HSC were index-sorted as HSC and scRNAseq analysis was undertaken to obtain the expression data of the sorted cells. Several clusters were defined for both BM-derived HSC and MPB-derived HSC. Single cell expression analysis of approx. 100 genes known to be specific for G0 (quiescent cells) or G1, S or G2M (cycling cells) revealed extensive expression of G0 specific genes in both BM and MPB derived. These results suggest that during cycling, the HSC choose G1 to follow the previous M phase, while they switch to G0 following their last cell division in BM.

## Results

We set out to compare human HSC from young adult BM with mobilized blood cells from 3 donors, 1 mobilized using only G-CSF and 2 using G-CSF plus Plerixafor. HSC were obtained using index-sorting as described in methods (Supplemental Figure 1A and 1B). Briefly, bone marrow was stained with CD34-APC-Cy7, CD38-APC, CD90-Alexa Fluor 488, CD45RA-BV786. The Lin mix contained antibodies directed against CD3, CD4, CD8, CD11b, CD14, CD19, CD20, CD56 and CD235a. Cells were index-sorted as HSC: Lin-CD34+ CD38-CD90+ CD45RA-; MPP: Lin-CD34+ CD38-CD90-CD45RA- or LMPP: Lin-CD34+ CD38-CD90-CD45RA+. The index sort data (supplemental figure 1B) confirms post sequence analysis that the cells used are all from within the sort gates. Supplemental Figure 2A shows the presence of the individual donors in the HSC clusters. Contributions are similar for the three MPB samples and show more variability for the four BM samples.

### Cell cycle analysis

We use scRNAseq data, performed as described^14,15^, to distinguish between the G0, G1, S and G2M phases, with special emphasis on the G0/G1 distinction. Normally, most HSC are in G0 (quiescent). Rapid entry into cycle may be essential to survive severe blood loss, but this also increases vulnerability to blood cancers^16^. The transition from G0 to G1 allows G1 phase repair of DNA damage and also coordinates DNA synthesis^17^. To distinguish the various cell cycle phases based on mRNA data, we defined a list of 96 genes known to be preferentially expressed in the various phases (Table 1 Supplemental). We exported single cell data as the fraction of cells expressing and average expression per cluster for each of the 42,505 transcripts analyzed. Overall, (supplemental Figure 2B) G1/S/G2/M expression is limited to BM clusters 3 and 4 and approx. 10% of the cells in cluster 5. MPB expresses G1/S/G2M in less than 10% of clusters 0-4 and in 100% of cluster 5, which contains few cells (1.3% of the BM cells). There are differences in the G0 genes expressed in BM and MPB, with *MPL, MLLT3* and *STAT6* expressed at higher levels in MPB (Supplemental Figure 2C).

Figure 1A shows the outcome in detail. HSC-gated cells sorted from BM and MPB show G1, S and G2M expression mainly in cluster 3 and 4 or cluster 5 respectively. The fraction expressing G0 specific genes tends to be lower in MPB than in BM. Analysis of HSPC shows similar results (Figure 1F). The absence of DNA repair gene expression (Supplemental Fig 4A) in the MPB HSC, other than in the dividing cells in cluster 5, confirms the G0, rather than G1, phase of the majority of MPB HSC.

**Figure 1.**
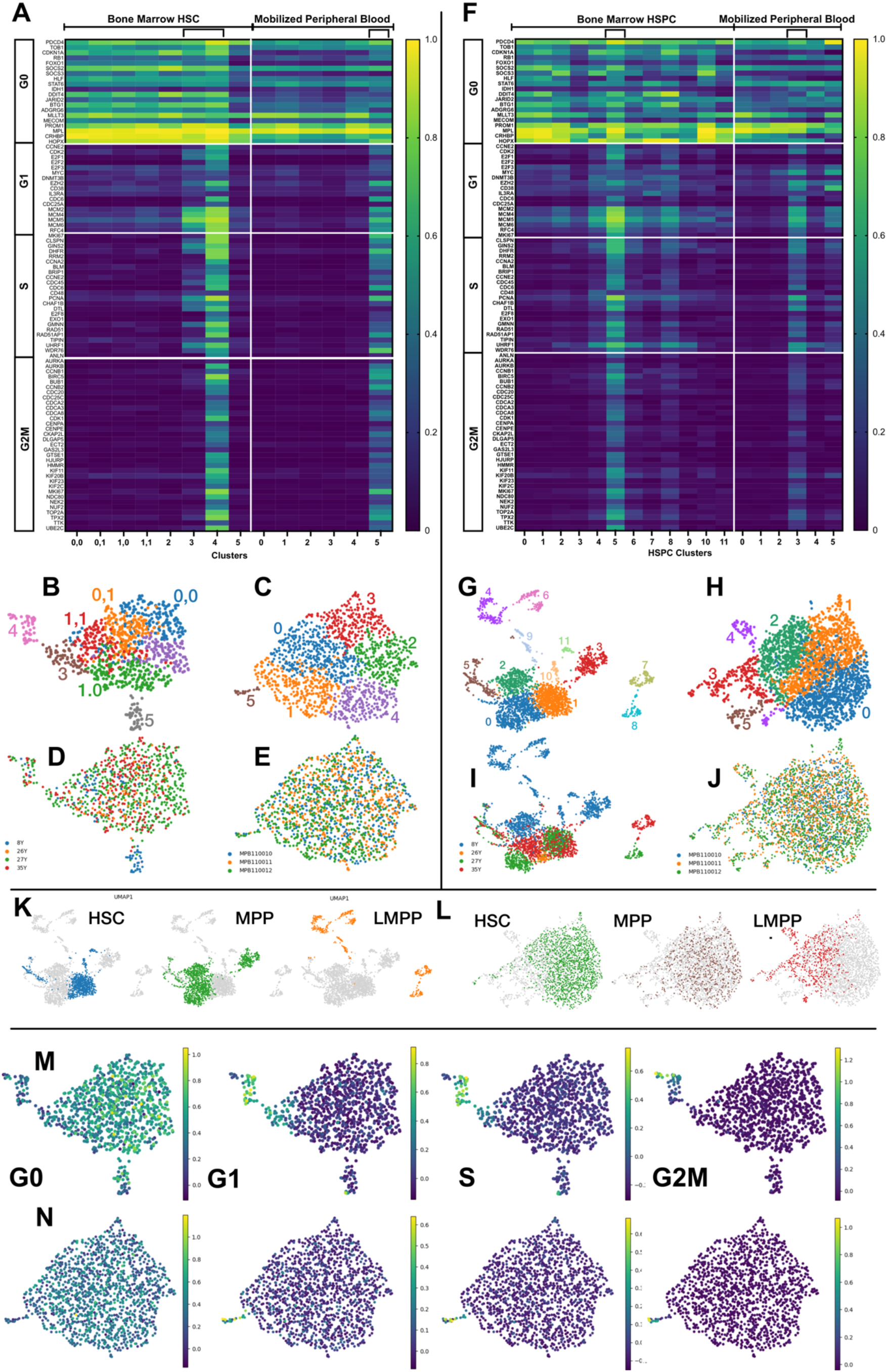
A-E and M, N show data gated as HSC, F-L data gated as HSPC (see supplemental data 1 for gating). (A) BM HSC expression of 94 genes (listed in supplemental table 1) selected to be expressed in different cell cycle phases. G0 genes are expressed in all clusters in BM and MPB. G1, S and G2M specific genes are only expressed in 1 or 2 clusters. Expression of the G0 fraction is higher in BM derived HSC. (B) shows the 6 clusters obtained through unsupervised clustering of HSC-gated BM donors. (C) shows similar cluster data for MPB. (D) and (E) shows the distribution of the different donors, and correlation with the clusters, in BM and MPB respectively. (F) shows similar data to (A) but the cells are gated as HSPC. (G) shows clustering data (12 clusters) for HSPC-gated BM. (H) shows the same with 6 MPB clusters. (I) and (J) show the distribution of the contributing donors to HSPC gated BM and MPB respectively. (K) shows the distinct distribution of HSC-gated, MPP-gated and LMPP-gated BM cells. (L) shows the same distribution for MPB HSC, MPP and LMPP. The distribution largely overlaps. (M) and (N) show scRNAseq analysis for BM (M) and MPB (N) respectively for the different sets of the 94-cell cycle phase specific genes as defined above. G0 specific genes are expressed throughout, while G1, S and especially G2M genes are limited. G0 cells are expressed at a higher level in (M) than (N).

Figure 1B-E (HSC gated) shows analysis for four BM and three MPB samples while Figure 1G-J shows the same for HSPC gated cells. (B) depicts the clusters for BM and (C) for MPB HSC. It also distinguishes donor identities (age or sampleID) per cluster (Figure 1D, E). Most contain cells from all donors. BM cluster 5 e.g. is mostly derived from one donor. This is more obvious for the HSPC clustering (Figure 1G - J). The figure also shows the clusters occupied by FACS groups (HSC, MPP, LMPP). BM HSPC clusters tend to have mostly well separated FACS groups (Figure 1K) while MPB HSPC FACS groups overlap (Figure (1L).

The figure also shows single cell G0/G1/S/G2M expression data for BM (M) and MPB HSC (N). Most cells express G0-specific genes while G1, S and G2M are expressed in a small subset. The main difference between BM and MPB is the expression level. Both express G0 genes in most cells. Expression is higher (yellow) in BM than in MPB (green). *MKI67* Expression confirms this: 7.4% of the BM HSC express *MKI67* while only 1.5% of the MPB HSC do.

Figure 2A compares expression in BM and MPB of approximately 200 transcription factors while Figure 2B shows this for signal transduction genes, both listed in supplemental data. The cell fraction that expresses these transcription factors tends to be higher for BM than for MPB, e.g. *PTEN, CDC42* and *RAC1*, all involved in HSC egress^18^. There are exceptions like *JUN*. To emphasize this panel, we show (Figure 2C) from the same dataset, the transcription factors that are expressed at a higher fraction and average levels in MPB (data shows MPB-BM as well as (Figure 2D) signal transduction genes. The same highly transcribed genes tend to stand out when showing fraction or average. For transcription factors *JUN*, part of the AP-1 complex that regulates proliferation and differentiation, albeit not HSC specific, stands out^19^, followed by *NFE2*, involved in controlling HSC self-renewal^20^, *ATF7*, a regulator of chromatin without a clear HSC function and *CDKN1B*, which promotes G1 arrest and HSC quiescence^21^. For the signal transduction genes this includes *CDC42*, a GTPase involved in HSC polarity and aging^18^, and *EGR1*, a transcription factor that regulates quiescence and niche anchoring^22^. (Figure 2E, F) shows the similar data but rather than a limited set of pre-selected genes it was calculated as MPB-BM for all 42,505 transcripts analyzed. The figure only shows the 50 genes with the highest– and lowest– values. Among the genes expressed at the highest levels in MPB over BM are various quiescence related genes such as *ARID1A*, a chromatin remodeler which is critical for maintaining the HSC pool size^23^, *HUWE1*, essential for a normal G0 pool size^24^, *DDX6*, essential for stemness and long-term reconstitution^25^ and *ANXA2*, which binds *CXCL12* and thus facilitates migration and adhesion^26^. Other genes that differ between MPB and BM include many histones and HLA genes as well as non-protein encoding RNA genes. Similar patterns are seen when comparing average expression rather than fraction expressing (Supplemental Figure 3A, B). Genes overexpressed in BM contain some, but not many, HLA genes. Our data shows that the switch to G0 is partial (lower-level expression) as the cells transition through the vasculature. Initial testing reveals no clear differences in expression between BM and MPB for genes expressed in (almost) all cells at high levels, such as CD74 (supplemental Fig. 3C) and several other genes such as *MALATH1, FOS, MPL* and *B2M*. Several genes are expressed at significantly higher levels in MPB than in BM, e.g. *CDC42* and *JUN*.

**Figure 2.**
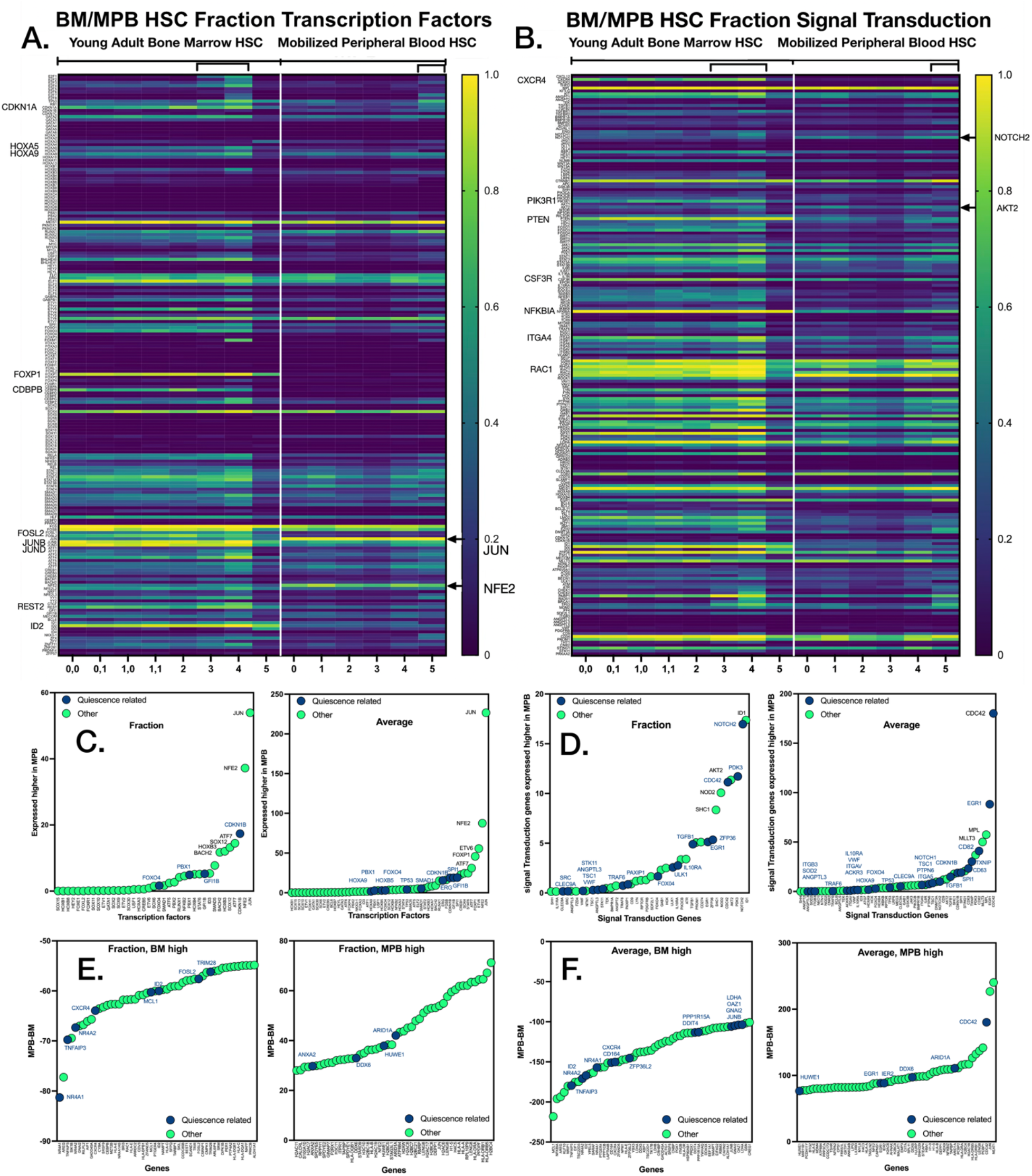
All data gated as HSC. (A) Heatmap showing the fraction of cells per cluster expressing each of approximately 200 transcription factors. (B) shows a heatmap depicting the fraction of cells per cluster expressing each of 200 signal transduction genes. Genes for A and B are listed in Supplemental Data. (C) shows the cell fraction and average expression of transcription factor genes (same as in Fig 2A) expressed at higher level in MPB. (D) shows the cell fraction and average expression level of signal transduction genes (same as in Supplemental Fig 2B) that are expressed at higher levels in MPB than in BM. JUN and NFE2 (transcription factors) and CDC42 and EGR1 (signal transduction factors) stand out as having the largest difference in expression. (E) shows the genes with the largest difference in fraction expressed of all genes present in the scRNAseq data (the 50 highest differentially expressed HSC genes in BM and in MPB are shown. Blue symbols show genes involved in HSC quiescence, green symbols show expression of other genes, in the right plot mostly histones and HLA genes. JUN, while not direct involved in HSC quiescence, like in (C) is expressed at clearly higher levels in MPB. (F) shows average expression data for BM and MPB. The plots show the 50 highest differentially expressed genes overexpressed in BM HS.

The release of HSC from the BM into circulation involves disruption of retention signals, esp. CXCR4 on HSC (Figure 2E). To see if this also involves other genes^27,28^, we investigated RNA expression of adhesion molecules (Supplemental Figure 4B). As expected, *CXCR4*, highly expressed in BM HSC, is not expressed in MPB HSC. Integrins, esp VLA4, i.e. α4 (ITGA4) and β1 (ITGB1), and VLA-5 (α5 and β2) are expressed less in MPB HSC.

## Discussion

We aim to better understand the seeming contradiction that rapidly cycling HSC engraft well^1,2,13^. As published, G0 cells engraft better long-term than G1 cells with 100% of G0 cells engrafting and 0% of G1 cells following injection of 50 HSC^1^. These previous experiments were performed in mice, which allows a more detailed analysis and the use of highly purified HSC. Both in mice and human HSC the percentage in S/G2/M is significantly lower in MPB than in untreated BM HSC^10^.

Can we distinguish between the G0 and G1 phases of the circulating HSC following mobilization? To address this, we performed scRNAseq analysis on HSC that were index sorted from BM or MPB. G0 marker genes are expressed in all clusters of both BM and MPB. Similar to what has been published^10^ we find fewer cycling cells present in the MPB clusters. The cycling BM clusters (#3 & 4), contain 11.5% of the analyzed cells while the cycling MPB cluster #5 contains only 1.3% of the analyzed cells. Figure 1M, N also illustrates this low level of dividing cells in MPB.

scRNAseq analysis allows us to obtain expression data that may be helpful in better understanding the pathways that are involved in the rapid transition from dividing cells (G1/S/G2M) in quiescent (G0) cells. Figure 1 shows that, like BM, MPB expresses many G0 genes, but in less cells and at lower levels. Few of the G0 genes are expressed at higher levels in MPB when compared to BM, but a significant number are expressed at higher levels in BM, e.g. HOPX and CRHBP. Our data shows that the switch to G0 is partial as the cells transition through the vasculature, suggesting an ongoing process. However, among the genes expressed at the highest levels in MPB over BM (Fig 2E) there are various quiescence related genes such as ARID1A^23^, DDX6^25^ and ANXA2^26^. They may hold the key to the mechanism behind this switch of G1 to G0.

Speculation. Our insights in this process are based on both human and mouse analysis: Human experiments are ethically limited. Mobilization starts with expansion of HSC in the bone marrow. We propose that it will be critical to explain how HSC in BM switch from the G1 program (continued cell divisions, poor engraftment capacity) to the G0 program (no more cell divisions, efficient engraftment). It seems that HSC “count” the number of cell divisions before switching to G0 and leaving the bone marrow. Elucidating the mechanisms behind this transition should allow for better control of expansion and engraftment. It has wider implications. In cancer, hypoxic conditions may force cells into G0, enabling them to evade the effects of chemotherapy^29^. A better understanding of the regulation of G0 and G1 could dramatically improve outcomes.

## Acknowledgements

Grants: this work was supported by NCI/NIH R35, 5R35CA220434, Ludwig and a CIRM Centers of Excellence grant awarded to Michael Snyder: Grant GC1R-06673-A, that supported the sequencing efforts. We thank Tejaswitha Naik for laboratory management, Cathy Carswell-Crumpton and the ISCBRM flow cytometry core for assistance with FACS, Norma Neff, CCbiohub for sample and library preparation and the Stanford Center for Genomics and Personalized Medicine (SCGPM) for sequencing support. We thank Matthew Porteus for making MPB samples available.

The datasets presented in this study can be found in online repositories.

