## Supplemental Files for "Human Haematopoietic Stem Cells released into circulation following mobilization express multiple G0-associated quiescence markers"

**Supplemental Data.**

### Methods

**Flow Cytometry: Analysis and Cell Sorting.** The following panel of antibodies (Caltag/Invitrogen and BD Biosciences) was used for analysis and sorting of human hematopoietic stem and progenitor populations: PE-Cy5–conjugated anti-human lineage markers (anti-CD3, S4.1; anti-CD4, S3.5; anti-CD8, 3B5; anti-CD11b, ICRF44; anti-CD14, TU.K.4; anti-CD19, SJ25-C1; anti-CD20, 13.6E12; anti-CD56, B159; anti-CD235A, GA-R2), CD34 (8G12) APCCY7 conjugated, CD38 (HIT2), PECy7 conjugated, CD45RA (MEM56) BV786 conjugated, CD90 (5E10), FITC conjugated and CD123 (6H6), PE-conjugated.

For analyses and sorting, except otherwise noted below, cells were stained with the appropriate antibody combinations for 30–60 min on ice, and dead cells were excluded by propidium iodide staining. Cells were analyzed and sorted using a FACSAria II cytometer (BD Biosciences). Analysis of flow cytometry data was performed using FlowJo Software (Becton and Dickinson).

**Single cell RNA sequencing (scRNA-seq).** This was performed as described[4–6]. Supplemental Figure 1a shows analysis for bone marrow (BM) derived HSC and shows this information for three MPB samples and four BM samples. Following normalization and removal of batch effect variation using ComBat [1], we applied the Leiden algorithm [2] and visualized the data with UMAP uniform manifold app [3] and non-supervised Leiden clustering. Cluster numbers range from 6 to 12 clusters per sample. Normalization did not include removal of ribosomal genes and non-protein coding RNA-transcripts. Analysis and clustering were done on 943 HSC from BM HSC samples in 8 clusters and 2823 BM HSPC samples in 12 clusters. As for MPB, analysis included 1437 HSC in 6 clusters and 2821 HSPC also in 6 clusters. Minor clusters (less than 10 cells) were excluded. We used sets of approximately 200 genes to test transcription factors, signal transduction, DNA repair and cell adhesion genes. There is some overlap between the groups, this doesn't affect the analysis.

**Tested transcription factor genes (Figure2A, C).** The transcription factors group contained the following genes: E2F1, E2F2, E2F3, E2F4, E2F5, E2F6, E2F7, E2F8, TP53, RB1, CDKN1A, CDKN1B, CDKN1C, GATA2, GATA3, GATA1, GATA4, GATA5, GATA6, HOXA1, HOXA2, HOXA3, HOXA4, HOXA5, HOXA7, HOXA9, HOXA10, HOXA11, HOXA13, HOXB1, HOXB2, HOXB3, HOXB4, HOXB5, HOXB6, HOXB7, HOXB8, HOXB9, HOXC4, HOXD1, HOXD3, HOXD4, HOXD8, HOXD9, PBX1, PBX2, PBX3, MEIS1, PNOX1, PNOX2, RUNX1, RUNX2, RUNX3, TAL1, GATA2, MYC, MYCN, MYCL, USF1, USF2, BHLHE40, BHLHE41, HEY1, HEY2, HEYL, FLI1, ERG, ELF1, ELF2, ELF3, ELF4, ELF5, GABPA, GABPB1, ETV1, ETV2, ETV3, ETV4, ETV5, ETV6, ETV7, SPI1, FOXO1, FOXO3, FOXM1, FOXA1, FOXA3, FOXC1, FOXC2, FOXE1 , FOXF1, FOXF2, FOXL1, FOXL2, , FOXP1, FOXP2, FOXP4, FOXR1, CEBPA, CEBPB, CEBPD, CEBPE, CEBPG, CEBPZ, SOX2, SOX17, SOX4, SOX5, SOX6, SOX7, SOX8, SOX9, SOX10, SOX11, SOX12, SOX12, SOX13, SOX15, SOX18, SOX21, SOX30, RELA, NFKB1, NFKB2, RELB, REL, STAT1, STAT2, STAT3, STAT5A, STAT5B, STAT6, STAT4, SMAD1, SMAD2, SMAD3, SMAD4, SMAD5, SMAD6, SMAD7, SMAD9, HLF, CEBPD, FOS, FOSB, FOSL1, FOSL2, JUN, JUNB, JUND, ATF1, ATF2, ATF3, ATF4, ATF5, ATF6, ATF7, CREB1, CREB3, CREB4, CREB5, BACH1, BACH2, NFE2, NFE2L2, NRF1, NFE2L3, YY1, CTCF, REST, GFI1, MECOM, BCL6, ID1, ID2, ID3, ID4, NKX3-1, ZFX, ZFY, ZNF711, ZNF281, PRDM14, ZFP57.

**Tested signal transduction genes (Figure 2B, D).** The signal transduction genes group, depicted in Fig 2B,D, includes: CXCL12, CXCR4, ACKR3, THPO, MPL, KITLG, KIT, ANGPT1, ANGPT2, TEK, TGFB1, TGFB2, TGFBR1, TGFBR2, BMPR1A, BMPR1B, BMPR2, ACVR1, ACVRL1, ENG, NOTCH1, NOTCH2, JAG1, JAG2, DLL1, DLL4, RBPJ, HES1, HEY1, NUMB, WNT3A, WNT5A, FZD4, FZD6, LRP5, LRP6, CTNNB1, APC, GSK3B, SHH, PIK3CA, PIK3CB, PIK3CD, PIK3R1, AKT1, AKT2, MTOR, RPTOR, RICTOR, PTEN, TSC1, TSC2, FOXO1, FOXO3, FOXO4, SIRT1, SIRT3, SIRT7, JAK1, JAK2, JAK3, TYK1, STAT1, STAT3, STAT5A, STAT5B, IL6R, IL6ST, IL11RA, IL7R, CSF3R, EPOR, IL10RA, SOCS1, SOCS3, SH2B3, NFKB1, RELA, RELB, CHUK, IKBKB,

NFKBIA, TLR2, TLR4, TLR9, MYD88, IRAK1, TRAF6, NOD1, NOD2, ITGA4, ITGB1, ITGA6, ITGA5, ITGAV, ITGB2, VCAM1, SELP, CD44, CD63, RHOA, RAC1, RAC2, CDC42, ROCK1, VAV1, SRC, LYN, FYN, HCK, SYK, PTPN6, PTPN11, SHC1, GRB2, GAB1, HIF1A, EPAS1, MEG3, TXNIP, PRDX2, SOD2, GPX1, PDK2, PDK3, LDHA, NFE2L2, GPRC5C, GPRC5A, ADGRG1, GPR123, ADRB3, HRH2, NEO1, NTN1, CLEC9A, CD82, CD274, SLAMF1, CD48, GATA2, MEIS1, HOXA9, HOXA10, HOXB4, EGR1, BCL6, BCL2, BCL2L11, KLF4, TAL1, LMO2, SPI1, IKZF1, BMI1, EZH2, DNMT3A, G0S2, CDKN1C, CDKN1B, ID1, ID2, ID3, ZFP36, FSTL1, MECOM, MLLT3, PAXIP1, ATP6V0A1, ATG5, ATG7, BECN1, ULK1, ATM, ATR, CHEK1, CHEK2, PARP1, RAD51, BRCA1, TP53, MDM2, PF4, SDF2L1, ITGB3, ANGPTL2, ANGPTL3, ANGPTL5, VWF, PDGFRB, CD9, CD34, PROM1, THY1, CNR2, STING1, STK11, PRKAA2.

**Tested DNA repair genes (Figure 4A Supplemental).** ATM, ATR, PRKDC, TP53, CHEK1, CHEK2, MDC1, H2AX, RNF8, RNF168, TP53BP1, UIMC1, BRCC3, ABRAXAS2, MRE11, RAD50, NBN, RIF1, TOPBP1, CLSPN, LIG4, XRCC4, NHEJ1, PAXX, POLM, POLL, APTX, TDP1, BRCA1, BRCA2, PALB2, RAD51, RAD51B, RAD51C, RAD51D, XRCC2, XRCC3, RAD52, RAD54L, RAD54B, BARD1, BRIP1, EME1, MUS81, SLX1A, SLX4, GEN1, RBBP8, EXO1, DNA2, BLM, WRN, RECQL, RECQL4, SMARCAL1, ZRANB3, FANCA, FANCB, FANCC, FANCD2, FANCE, FANCF, FANCG, FANCI, FANCL, FANCM, ERCC4, FAN1, UBE2T, RFWD3, MSH2, MSH3, MSH6, MLH1, PMS2, PMS1, MLH3, PCNA, RFC1, MPG, TDG, UNG, MBD3, OGG1, MUTYH, MTHL1, SMUG1, APEX2, PARP2, XRCC1, LIG1, PNKP1, POLB, POLG, NEIL1, NEIL2, NEIL3, XPA, POLBB, POLG2, NEIL1, NEIL2, NEIL3, XPA, XPC, RAD23B, DDB1, DDB2, ERCC1, ERCC2, ERCC3, ERCC4, ERCC5, ERCC6, ERCC8, GTF2H1, GTF2H2, GTF2H3, GTF2H4, GTF2H5, ALDH2, ADH5, SPRTN, DCLRE1A, DCLRE1C, EME2, USP1, WDR48, TERT, TERC, POT1, ACD, TERF1, TERF2, TERF2IP, TINF2, RTEL1, CDKN1B, GADD45G, HUS1, RAD1, RAD9A, BCL2L1, BAX, PPM1D, CHD2, CHD4, SMARCA5, SMARCAD1, INO80, SRCAP, KAT5, EP300, CREBBP, SETD2, TIMELESS, TIPIN, RAD17, RFC2, SMC1A, SMC3, ESCO2, LIG3, APLF, RPA1, RPA2,

RPA3, SMC5, SMC6, SLFN11, TREX1, TREX2, RNASEH1, RNASEH2A, RNASEH2B, RNASEH2C

**Tested cell adhesion genes (Figure 4B Supplemental).** ITGA4, ITGB1, ITGA5, ITGAL, ITGB2, ITGA6, ITGA7, ITGA9, ITGAV, ITGB3, ITGA2B, ITGA2, ITGAM, ITGAX, ITGAE, ITGB7, ITGB5, ITGB8, ITGA1, ITGA3, ITGA8, ITGA10, ITGA11, ITGB4, ITGB6, SELE, SELP, SELL, SELPLG, CD44, GLG1, PODXL, CDH2, CDH11, CDH1, CDH5, CDH13, PCDH7, FAT1, FAT4, CDH6, CDH9, ALCAM, ESAM, F11R, JAM2, JAM3, ICAM1, ICAM2, ICAM3, ICAM4, ICAM5, VCAM1, NCAM1, PECAM1, CD99, SLAMF1, CD48, CD244, CD84, LY9, SLAMF6, SLAMF7, SIGLEC1, SIGLEC5, SIGLEC7, SIGLEC9, L1CAM, MCAM, CEACAM1, CEACAM6, CD47, SIRPA, SIRPB1, CD200, CD200R1, CD96, CRTAM, NECTIN2, NECTIN3, PVR, CADM1, IGSF11, VCAN, CNTN2, CNTNAP1, ROBO4, HMMR, THBS1, THBS2, SPARC, TNC, LAMA1, LAMA2, LAMA4, LAMB1, LAMC1, FN1, VTN, COL1A1, COL1A2, COL2A1, COL3A1, COL4A1, COL4A2, COL5A1, COL6A1, NID1, NID2, HSPG2, AGRN, OPN, POSTN, TGFB1, LGALS1, LGALS3, VCAN, CSPG4, BGN, DCN, LUM, NOTCH1, NOTCH2, NOTCH3, NOTCH4, JAG1, JAG2, DLL1, DLL3, DLL4, EFNB1, EFNB2, EFNA1, EFNA2, EPHB4, EPHB2, EPHA4, CXCR4, CXCL12, ACKR3, CCR1, CCR2, CCR5, CCR7, CXCR2, CX3CR1, ACKR1, CD9, CD81, CD82, CD151, TSPAN7, TSPAN32, TSPAN33, TSPAN18, NT5E, THY1, ENG, PDGFRA, PDGFRB, CD34, CD93, CD109, TNFSF9, MADCAM1, CD36, CD163, S1PR1, S1PR3, PLXNB2, PLXND1, NRP1, NRP2, ROBO1, SLIT2, PTPRC, CD24, CD52, CD99L2, BST2, GPIBA, GP6, CLEC9A, CLEC12A, SIGLEC15, EMILIN1.

|  | Gene | Phase | Description | DOI |
|---|---|---|---|---|
| 1 | PDCD4 | G0 | Plays a critical regulatory role in HSC maintenance | 10.3324/haematol.2019.236927 |
| 2 | TOB1 | G0 | T cell, HSC quiescence | 10.1111/cen3.12125 |
| 3 | CDKN1A | G0 | Cell cycle inhibitor, promotes quiescence | 10.4161/cc.25416 |
| 4 | RB1 | G0 | Enforces cell cycle exit and maintenance | 10.1016/j.cell.2007.03.055 |
| 5 | FOXO1 | G0 | Forkead TF, high in HSC, promotes self renewal. Induces cell cycle inhibitor genes | 10.3389/fimmu.2019.01016 |
| 6 | SOCS2 | G0 | Highly expressed in HSC, inhibitor in JAK/STAT | 10.1002/JLB.1A1217-480R |
| 7 | SOCS3 | G0 | Blocks JAK/STAT signaling, induces receptor degradation | 10.1084/jem.20160719 |
| 8 | HLF | G0 | Crucial for maintaining the self-renewal/multipotency of HSCs | 10.1038/s41586-022-04571-x |
| 9 | STAT6 | G0 | deficiency leads to increased numbers of committed myeloid progenitors | 10.1189/jlb.0903440 |
| 10 | IDH1 | G0 | Hypoxia response, critical for HSC quiescence, survival, and regenerative capacity | 10.3892/ol.2019.10256 |
| 11 | DDIT4 | G0 | Negative regulator of mTORC1, controls cell growth, proliferation, and metabolism | 10.1002/stem.2028 |
| 12 | JARID2 | G0 | Epigenetic regulator of HSPC. Balances quiescence, self-renewal, and differentiation, | 10.1182/blood-2014-10-603969 |
| 13 | BTG1 | G0 | negatively regulates cell cycle progression, acts as a tumor suppressor | 10.5483/BMBRep.2022.55.8.092 |
| 14 | ADGRG6 | G0 | adhesion G protein-coupled receptor, marks immature HSPCs | 10.1093/jbmr/zjae144 |
| 15 | MLLT3 | G0 | Epigenetic regulator, crucial regulator of HSC self-renewal and maintenance. | 10.1038/s41586-019-1790-2 |
| 16 | MECOM | G0 | Transcription factor required for HSC self-renewal and maintenance. | 10.1038/s41586-019-1790-2 |
| 17 | PROM1 | G0 | Cell surface marker of HSCs and progenitors, influencing activation and proliferation. | 10.1038/s41392-022-01301-7 |
| 18 | MPL | G0 | TPO receptor, indispensable for HSC maintenance, megakaryocyte differentiation | 10.1073/pnas.2017849118 |
| 19 | CRHBP | G0 | May contribute to regulating stem cell proliferation or differentiation, precise role unclear | 10.1038/s41467-025-57096-y |
| 20 | HOPX | G0 | Helps maintain HSC quiescence by regulating the CXCL12-CXCR4 signaling axis | 10.1038/s41388-020-1340-2 |
| 21 | CCNE2 | G1 | CyclinE2, G1/S transition | 10.1073/pnas.222491799 |
| 22 | CDK2 | G1 | Cyclin-dependent kinase 2, essential for G1/S transition | 10.1038/s42003-025-09038-z |
| 23 | E2F1 | G1 | Regulates cell cycle progression and HSC function. | 10.1038/sj.emboj.7600459 |
| 24 | E2F2 | G1 | Plays essential and partially redundant roles with E2F1 | 10.1128/MCB.23.10.3607-3622.2003 |
| 25 | E2F3 | G1 | E2F3a is highly expressed at the G1/S transition and regulated by E2F and Myc | 10.1128/MCB.02147-05 |
| 26 | MYC | G1 | Induction of critical cell cycle genes | 10.1016/j.stem.2020.09.004 |
| 27 | DNMT3B | G1 | DNA methyltransferase, highly expressed in long-term repopulating HSCs | 10.1126/sciadv.adu8116 |
| 28 | EZH2 | G1 | maintains the balance between HSC proliferation and differentiation by repressing differentiation | 10.7150/thno.53170 |
| 29 | CD38 | G1 | Distinguishes two populations of long-term (LT) HSCs. | 10.1371/journal.pbio.3002517 |
| 30 | IL3RA | G1 | Stimulates HSC exit from quiescence (G0 phase) | 10.1038/s41467-025-57096-y |
| 31 | CDC6 | G1 | Key regulator of DNA replication and cell cycle progression. | 10.1073/pnas.94.11.5611 |
| 32 | CDC25A | G1 | Cell cycle phosphatase that promotes progression from G1 to S. | 10.1038/s41698-025-00862-4 |
| 33 | MCM2 | G1 | Replicative helicase essential for DNA replication initiation and elongation. | 10.7554/eLife.80917 |
| 34 | MCM4 | G1 | Essential for the initiation and progression of DNA replication. | 10.1186/s12885-021-08344-z |
| 35 | MCM5 | G1 | Replicative helicase required for DNA replication initiation and progression during the S phase. | 10.14670/HH-24.299 |
| 36 | MCM6 | G1 | A crucial role in the initiation and elongation phases of eukaryotic DNA replication. | 10.1186/s12885-021-08344-z |
| 37 | RFC4 | G1 | Plays a crucial role in DNA replication and repair by loading the PCNA clamp onto DNA | 10.1186/s12885-021-08344-z |
| 38 | MKI67 | G1 | An important marker and regulator of active cycling in HSCs | 10.1186/s13287-023-03377-6 |
| 39 | CLSPN | S | Claspin, crucial cell cycle regulatory protein involved in DNA replication | 10.1111/febs.14594 |
| 40 | GINS2 | S | Promotes cell proliferation and inhibitits apoptosis. | 10.3892/ol.2018.8944 |
| 41 | DHFR | S | Dihydrofolate reductase role in the HSC cycle by supporting DNA synthesis | 10.1073/pnas.77.9.5140 |
| 42 | RRM2 | S | Catalyzes the rate-limiting step in deoxyribonucleotide production | 10.1016/j.gendis.2022.11.022. |
| 43 | CCNA2 | S | Cell cycle regulator that drives progression through the S phase and into mitosis | 10.1002/ajh.23952 |
| 44 | BLM | S | DNA helicase, crucial role in maintaining genomic stability, particularly in HSCs | 10.1046/j.1365-2249.1999.01060.x |
| 45 | BRIP1 | S | DNA helicase involved in DNA repair pathways. | 10.1371/journal.pgen.1011175 |
| 46 | CCNE2 | S | Key regulator of the cell cycle, G1/S phase transition | 10.1128/MCB.19.1.612. |
| 47 | CDC45 | S | Unwinds DNA at replication forks during the S phase | 10.1111/cpr.13257 |
| 48 | CDC6 | S | Crucial regulator of DNA replication initiation and cell cycle progression in HSCs | 10.1038/sj.embor.7400624 |
| 49 | NFE2L2 | S | Regulates migration and retention of HSCs in their bone marrow niches | 10.1128/MCB.00086-17 |
| 50 | CCN1 | S | Regulate stem cell niche interactions, promoting quiescence or activation. | 10.1084/jem.20050967 |
| 51 | CD48 | S | CD48 is mostly on short-term progenitors and ST-HSC. | 10.3389/fphys.2022.1009160 |
| 52 | PCNA | S | Sliding clamp for DNA polymerase during S phase, crucial for DNA synthesis and repair. | 10.1016/0145-2126(91)90101-x |
| 53 | CHAF1B | S | Critical role in DNA replication-linked S-phase chromatin assembly. | 10.1084/jem.20050967 |
| 54 | DTL | S | E3 ubiquitin ligase complex that targets proteins and prevents DNA re-replication | 10.1084/jem.20050967 |
| 55 | E2F8 | S | Transcriptional repressor, expressed during the S and G2 phases of the cell cycle | 10.4103/cjop.CJOP-D-22-00142 |
| 56 | EXO1 | S | Mismatch, double-strand break, excision repair (NER), and stalled replication forks | 10.1091/mbc.02-02-0030 |
| 57 | GMNN | S | Prevents DNA re-replication during the S phase | 10.1091/mbc.02-02-0030 |
| 58 | RAD51 | S | RAD51 repairing DNA double-strand breaks | 10.1091/mbc.02-02-0030 |
| 59 | RAD51AP1 | S | Crucial role in homologous recombination (HR) DNA repair by stimulating RAD51 activity | 10.1091/mbc.02-02-0030 |
| 60 | TIPIN | S | Critical regulator of the DNA replication fork and cell cycle | 10.1091/mbc.02-02-0030 |
| 61 | UHRF1 | S | Controls the establishment of DNA methylation patterns specifically during HSC division | 10.1091/mbc.02-02-0030 |
| 62 | WDR76 | S | Regulator of stem cell proliferation through controlling HSC cell cycle entry | 10.1091/mbc.02-02-0030 |
| 63 | ANLN | G2M | ANILLIN involved in cytokineses and cell cycle regulation | 10.1093/jmcb/mjy063 |
| 64 | AURKA | G2M | highly during the G2/M phase, orchestrates proper chromosome segregation | 10.1182/blood-2014-12-615401 |
| 65 | AURKB | G2M | Involved in chromosome alignment, segregation, and cytokinesis. | 10.3324/haematol.2011.054668 |
| 66 | CCNB1 | G2M | Cyclin B1 is a regulatory cell cycle protein with a critical role during the G2/M transition | 10.1093/jmcb/mjy063 |
| 67 | BIRC5 | G2M | SURVIVIN regulates cell division and prevents apoptosis | 10.1093/jmcb/mjy063 |
| 68 | BUB1 | G2M | BUB1 monitors kinetochore-microtubule attachments during mitosis | 10.1093/jmcb/mjy063 |
| 69 | CCNB2 | G2M | B-type cyclin, binds CDKs, important for G2M transitions | 10.1093/jmcb/mjy063 |
| 70 | CDC20 | G2M | Key protein in mitosis. E3 ubiquitin ligase complex that initiates chromosome separation | 10.1084/jem.20050967 |
| 71 | CDC25C | G2M | Critical regulatory phosphatase in the cell cycle, particularly at the G2/M transition. | 10.3892/ol.2015.2842 |
| 72 | CDCA2 | G2M | Recruits PP1 to chromatin, which is essential for chromatin decondensation. | 10.1093/jmcb/mjy063 |
| 73 | CDCA3 | G2M | Functionsas a trigger for mitotic entry. mediates ubiquitination and degradation of WEE1 | 10.1093/jmcb/mjy063 |
| 74 | CDCA8 | G2M | Plays a role in mitotic chromosome alignment, segregation, and spindle stabilization | 10.1093/jmcb/mjy063 |
| 75 | CDK1 | G2M | Governs mitotic entry and progression through the G2/M | 10.1093/jmcb/mjy063 |
| 76 | CENPA | G2M | Histone H3 variant that is specifically localized at centromeres | 10.1093/jmcb/mjy063 |
| 77 | CENPE | G2M | Kinesin-like motor protein that plays a key role in chromosome congression | 10.1093/jmcb/mjy063 |
| 78 | CKAP2L | G2M | A mitotic spindle protein with a key role in G2/M cell cycle regulation. | 10.1002/2211-5463.13864 |
| 79 | DLGAP5 | G2M | Key role in cell cycle regulation, during mitosis. | 10.1039/d4cb00155a |
| 80 | ECT2 | G2M | A guanine nucleotide exchange factor, critical role in cytokinesis | 10.1016/j.jbc.2021.101036 |
| 81 | GAS2L3 | G2M | A cytoskeleton orchestratorcrucial role in cytokinesis. GAS: growth arrests specific | 10.1074/jbc.M111.242263 |
| 82 | GTSE1 | G2M | Cell cycle-regulated protein primarily expressed during the S and G2/M phases. mitosis | 10.7554/eLife.101075 |
| 83 | HJURP | G2M | Critical histone chaperone involved in centromere function and chromosome segregation | 10.1016/j.cell.2009.02.040 |
| 84 | HMMR | G2M | Microtubule-associated protein. participates in mitotic spindle dynamics | 10.3389/fphar.2024.1361424 |
| 85 | KIF11 | G2M | Kinesin motor protein essential for mitotic spindle assembly and chromosome segregation. | 10.1016/j.yexcr.2024.113975 |
| 86 | KIF20B | G2M | A plus-end-directed motor protein involved in mitotic cell division, specifically in cytokinesis | 10.1089/mab.2019.0016 |
| 87 | KIF23 | G2M | Mitotic motor protein essential for cytokinesis, spindle midbody formation during cell division. | 10.1038/s44318-024-00327-7 |
| 88 | KIF2C | G2M | Crucial role in cytokinesis and mitotic spindle formation, mitotic progression | 10.1093/nar/gkaf476 |
| 89 | MKI67 | G2M | An important marker and regulator of active cycling in HSCs | 10.1186/s13287-023-03377-6 |
| 90 | NDC80 | G2M | Core component of the kinetochore complex, critical for accurate chromosome segregation. | 10.7150/jca.96070 |
| 91 | NEK2 | G2M | S/T kinase that plays a pivotal role in regulating the cell cycle, particularly during mitosis. | 10.1016/j.ejcb.2012.03.009 |
| 92 | NUF2 | G2M | Critical kinetochore protein, essential for chromosome alignment, segregation, cell division | 10.7150/ijbs.80737 |
| 93 | TOP2A | G2M | Critical enzyme involved in DNA topology modulation during the cell cycle | 10.1155/2021/4092635 |
| 94 | TPX2 | G2M | Microtubule-associated protein, regulates mitotic spindle assembly and mitosis. | 10.1002/jcb.30205 |
| 95 | TTK | G2M | Essential role in the spindle assembly checkpoint (SAC) during mitosis | 10.1002/ctm2.1544 |
| 96 | UBE2C | G2M | crucial in the ubiquitin-proteasome system regulating cell cycle progression | 10.1016/j.dnarep.2025.103901 |

Table 1 Supplemental

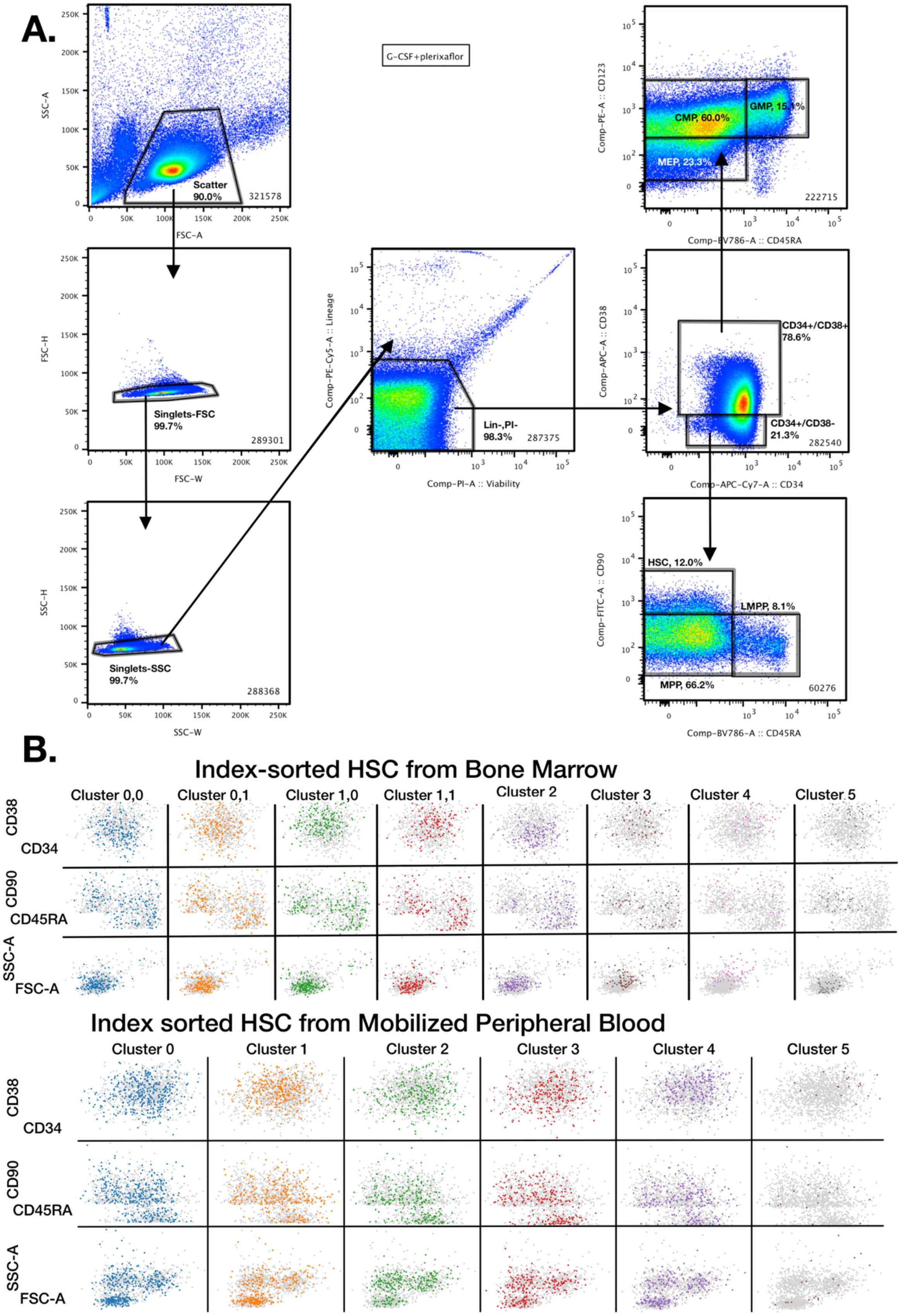
A.
G-CSF+plerixaflor
Scatter 90.0%
321578
SSC-A
FSC-A
Singlets-FSC 99.7%
289301
FSC-H
FSC-W
Singlets-SSC 99.7%
288368
SSC-H
SSC-W
Lin-,PI- 98.3%
287375
Comp-PE-Cy5-A :: Lineage
Comp-PI-A :: Viability
CMP, 60.0%
GMP, 15.1%
MEP, 23.3%
222715
Comp-PE-A :: CD123
Comp-BV786-A :: CD45RA
CD34+/CD38+ 78.6%
CD34+/CD38- 21.3%
282540
Comp-APC-A :: CD38
Comp-APC-Cy7-A :: CD34
HSC, 12.0%
LMPP, 8.1%
MPP, 66.2%
60276
Comp-FITC-A :: CD90
Comp-BV786-A :: CD45RA
B.
Index-sorted HSC from Bone Marrow
Cluster 0,0
Cluster 0,1
Cluster 1,0
Cluster 1,1
Cluster 2
Cluster 3
Cluster 4
Cluster 5
CD38
CD34
CD90
CD45RA
SSC-A
FSC-A
Index sorted HSC from Mobilized Peripheral Blood
Cluster 0
Cluster 1
Cluster 2
Cluster 3
Cluster 4
Cluster 5
CD38
CD34
CD90
CD45RA
SSC-A
FSC-A

Figure 1 Supplemental

***Figure 1 supplemental. Surface markers of the MPB cells HSPC.*** *Gating of the HSC and MPP/LMPP as used in this study. (A) The gating path used to sort HSC. The shown example is from CD34-enriched mobilized peripheral blood. HSC as used are Lin-CD34+CD38-CD90+CD45RA-. (B) shows the HSC index sort data from BM (top) and MPB (bottom) and confirms that the cells in the different clusters are part of the sort data. HSC in BM clusters tend to be part of the sorted cells, eg cluster 2 has a lower CD38 staining whereas cluster 1 is higher. Larger cells (by FSC/SSC) are present in the clusters expressing cell division markers, clusters 3 and 4 BM and cluster 5 MPB. MPB HSC are more evenly distributed.*

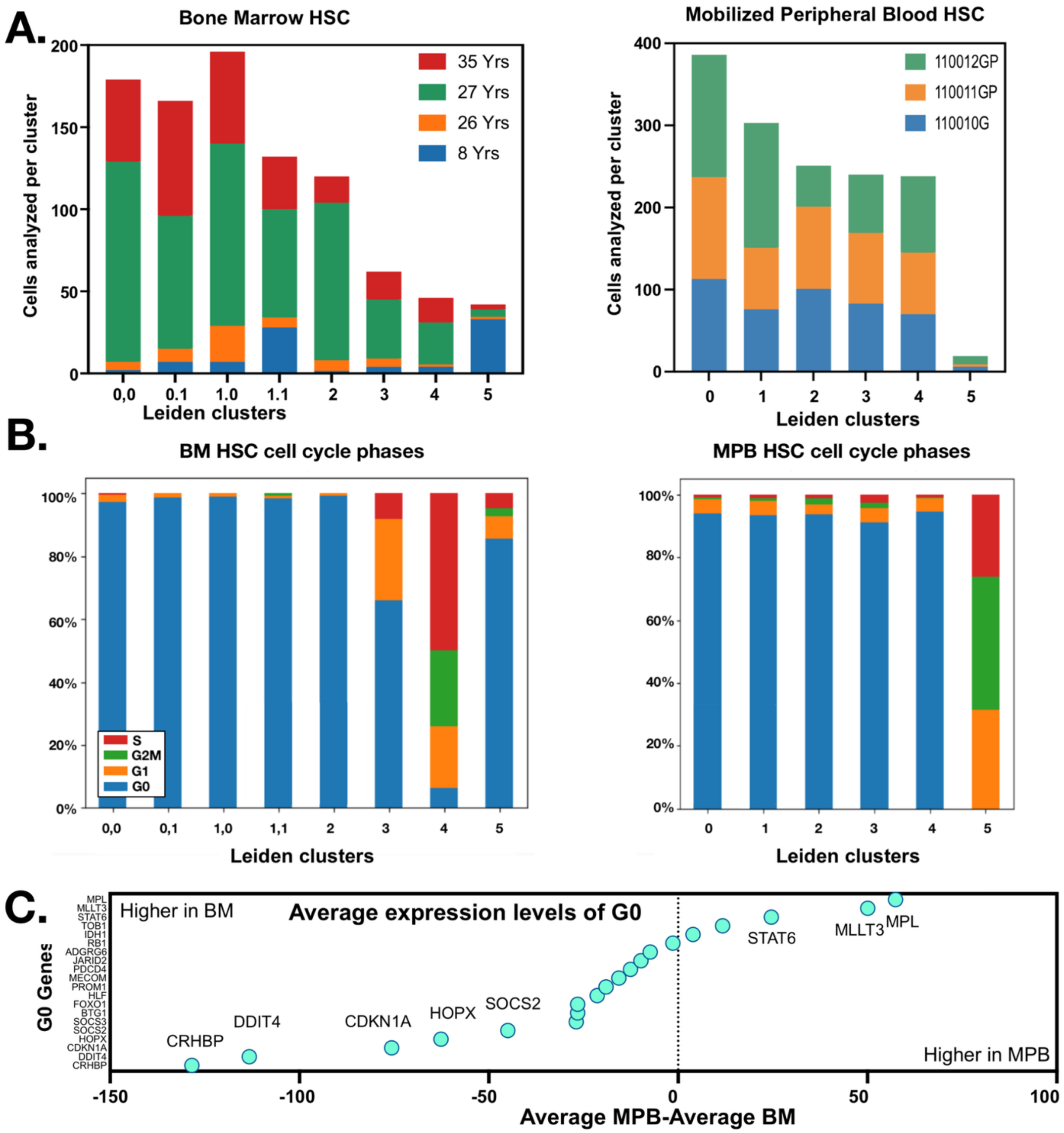


***Figure 2 supplemental. Cluster composition and G0 genes.*** *(A) The bargraphs show the contribution of the different donors to each of the clusters of BM and MPB HSC. (B) This panel shows the percentage of BM- and MPB-derived HSC in each cluster in the various cell phases (G0, G1, S and G2M). (C) The average expression level of G0 genes in MPB and BM. MPL and MLLT3 show the highest specific expression in MPB. The fraction of cells expressing these genes is high too (Figure 1A)*

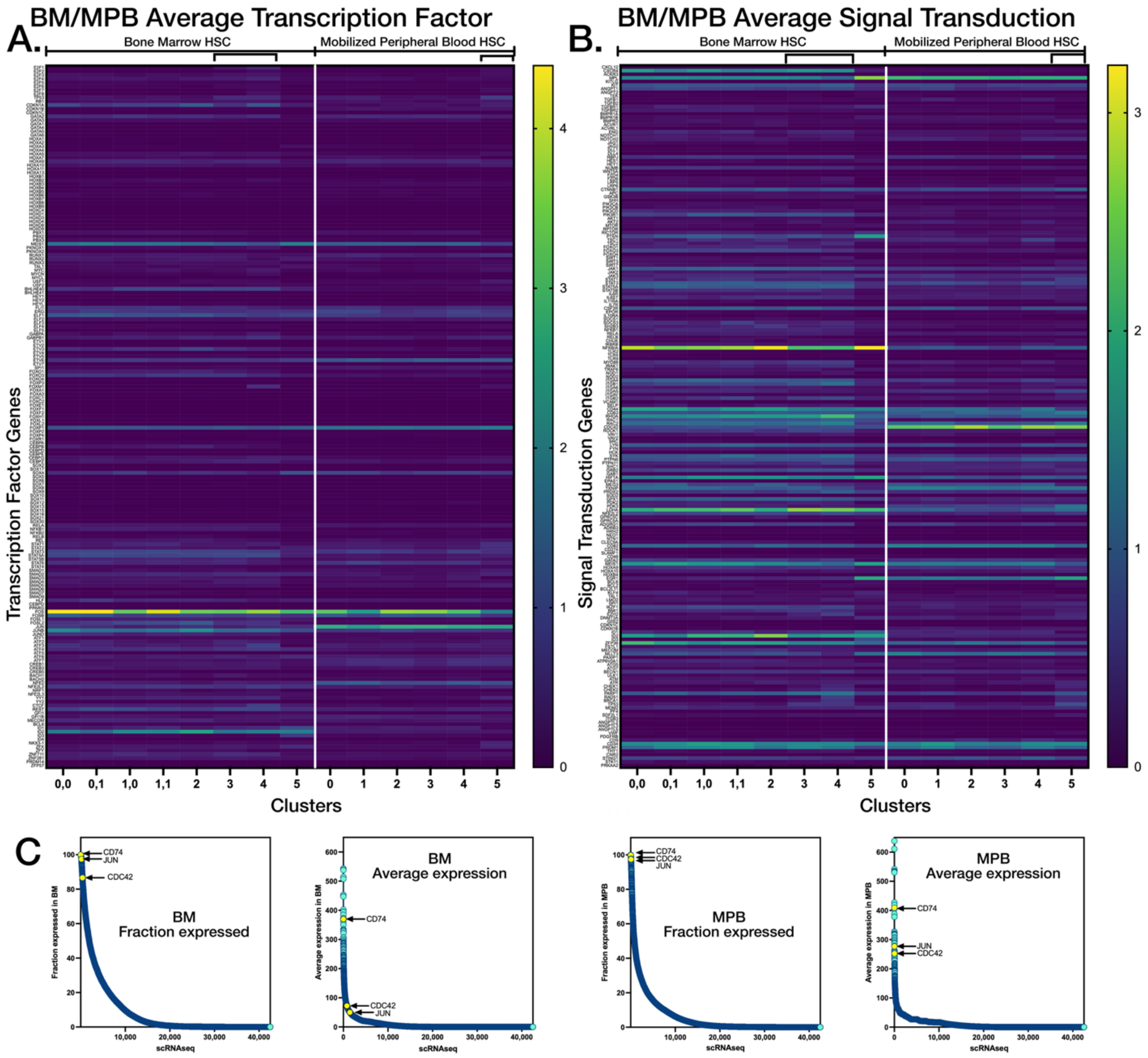


***Figure 3 supplemental. Expression of transcription factor and signal transduction genes.*** *Average expression per cluster. Data gated as HSC. (A) The heatmap shows the average expression per cluster for 200 transcription factor genes, as listed in the supplemental data. (B) shows a similar heatmap, depicting average expression for 200 signal transduction genes. (C)The bottom plots show the fraction of cells expressing- and average expression per cell-for BM and MPB HSC. While CD74, JUN and CDC42 are expressed in almost all BM and MPB HSC, average expression per cell of JUN and CDC42 is higher in MPB.*

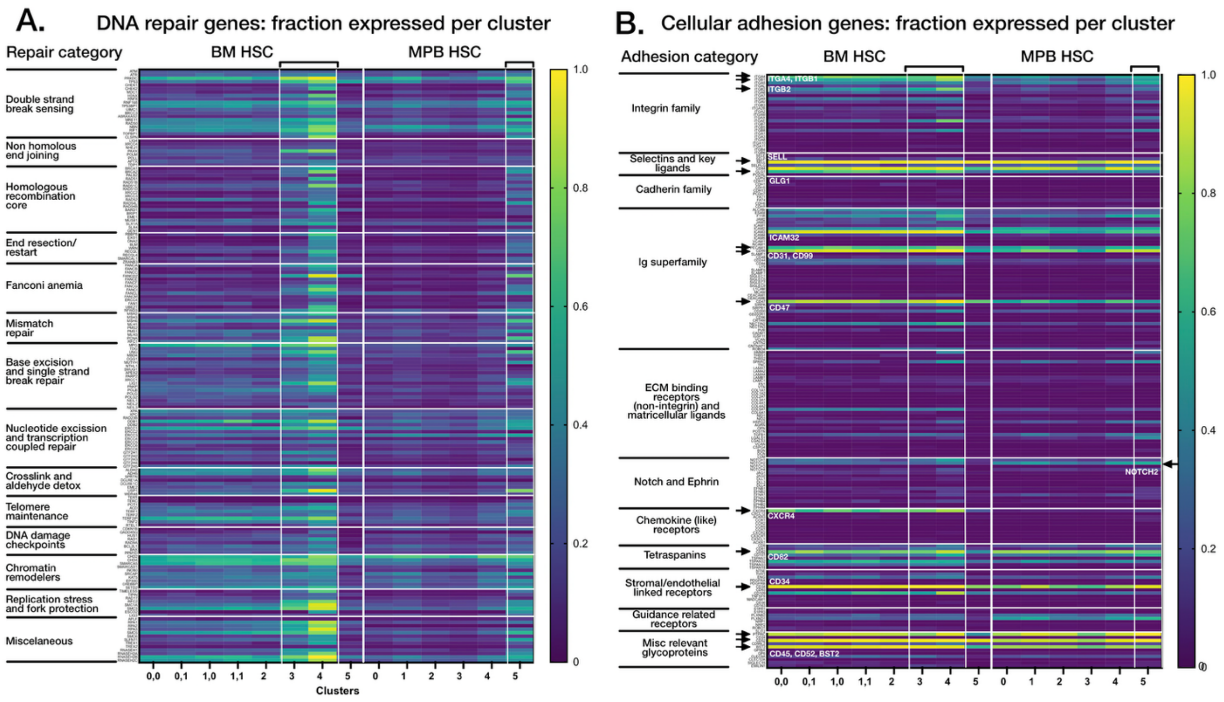


***Figure 4 supplemental. Expression of DNA repair genes and cellular adhesion genes in HSC.*** *(A) Fraction of HSC expressing DNA repair genes per cluster ranging from 1 (all cells express this gene) to 0 (none do). DNA repair genes are mostly expressed in dividing cell clusters. This is not different for MPB, confirming that these cells are not in G1. (B) Fraction of the HSC expressing cellular adhesion molecules. MPB derived cells express lower levels of integrin family members, and unlike BM, express no CXCR4. Integrin α4 and β1 are expressed at lower levels in MPB HSC. Both: Data gated as HSC. Dividing BM HSC are present in clusters 3 and 4. Dividing MPB HSC are present in cluster 5. The genes depicted in this figure are listed in the supplemental data section.*
